# Proteomics analysis of extracellular matrix remodeling during zebrafish heart regeneration

**DOI:** 10.1101/588251

**Authors:** Anna Garcia-Puig, Jose Luis Mosquera, Senda Jiménez-Delgado, Cristina García-Pastor, Ignasi Jorba, Daniel Navajas, Francesc Canals, Angel Raya

## Abstract

Adult zebrafish, in contrast to mammals, are able to regenerate their hearts in response to injury or experimental amputation. Our understanding of the cellular and molecular bases that underlie this process, although fragmentary, has increased significantly over the last years. However, the role of the extracellular matrix (ECM) during zebrafish heart regeneration has been comparatively rarely explored. Here, we set out to characterize the ECM protein composition in adult zebrafish hearts, and whether it changed during the regenerative response. For this purpose, we first established a decellularization protocol of adult zebrafish ventricles that significantly enriched the yield of ECM proteins. We then performed proteomic analyses of decellularized control hearts and at different times of regeneration. Our results show a dynamic change in ECM protein composition, most evident at the earliest (7 days post-amputation) time-point analyzed. Regeneration associated with sharp increases in specific ECM proteins, and with an overall decrease in collagens and cytoskeletal proteins. We finally tested by atomic force microscopy that the changes in ECM composition translated to decreased ECM stiffness. Our cumulative results identify changes in the protein composition and mechanical properties of the zebrafish heart ECM during regeneration.

## Introduction

According to the World Health Organization, 1/3 of all global deaths are due to cardiovascular diseases (1), which makes them the leading cause of morbidity and mortality among humans (2). This is generally thought to be the consequence of the limited capacity of the adult mammalian heart to recover after injury. Although some studies have reported that mammalian cardiomyocytes can experience limited proliferation after cellular damage, this is not sufficient to completely recover the lost tissue (3–5). After a lesion, the damaged cardiac tissue is replaced by a fibrotic scar, which reduces heart performance and may eventually lead to heart failure (6, 7).

Regeneration is a process by which organisms restore organs or tissues lost to injury or experimental amputation. This complex process gives rise to a tissue or organ nearly identical to the undamaged one. The understanding of regenerative processes is expected to help designing regenerative medicine strategies to address a variety of diseases. Even though neonatal mice have been shown to regenerate their hearts after an injury (8–13), the par excellence regenerative organisms are non-mammalian vertebrates such as urodele amphibians and zebrafish (14–17). Zebrafish can regenerate their heart after a 20% amputation (18, 19), cryoinjury (20–23) or genetic ablation(24). It has been demonstrated that this process occurs by limited dedifferentiation and proliferation of cardiomyocytes through the expression of cell-cycle regulators such as *plk1* and *mps1*, or chromatin-remodeling factors such as *brg1* (18, 25). Not only cardiomyocyte proliferation is essential for cardiac regeneration, but also cell migration, with cardiomyocytes near to the damaged tissue contributing to the regenerated tissue (26, 27). Moreover epicardium activation and migration are also required for the completion of the process (28–30). In contrast with our understanding of the cellular bases underlying zebrafish heart regeneration, the role of the extracellular matrix (ECM) in this process has received little attention (31–33).

Collagens, proteoglycans and glycoproteins form a complex structure, the ECM, which provides structural and mechanical support to tissues. In addition to a structural role, the ECM also instructs neighboring cells through biochemical and biomechanical signals that result in distinct biological responses (34, 35). Indeed, the composition and mechanical properties of the ECM play crucial roles determining cell behaviors such as proliferation, migration, differentiation, and apoptosis (36, 37). ECM remodeling is important during development, wound healing and disease states like cancer (36, 38–41). Studies using artificial ECM scaffolds have showed that physical properties of the ECM such as pore size, stiffness, fiber diameter, and chemical crosslinking also have effects on cell fate (42, 43). For instance, substrate stiffness induced myoskeleton reorganization, thus determining the cell shape of rat and mice neonatal cardiomyocytes, whereas dedifferentiation of neonatal cardiomyocytes and enhanced proliferation occurred in compliant matrices (43).

The role of the ECM during zebrafish heart regeneration has received little attention thus far. Gene expression analysis performed in regenerating hearts identified transcripts encoding ECM-related proteins among the most differentially expressed during this process (44–46). These included inhibitors of metalloproteinases, matrix metalloproteinases, and tenascin C (tnc), *tnc* transcripts being found overexpressed in the border zone between the healthy myocardium and the injury site (21, 47). Another study showed that fibronectin (fn) synthesis by epicardial cells was essential for zebrafish heart regeneration, probably by providing cues for cardiomyocyte migration (31). A role of hyaluronic acid (HA) signaling during zebrafish heart regeneration has also been proposed, because components of this pathway were found expressed in response to injury, and blocking HA signaling impaired regeneration (32). These three ECM components (tnc, fn, and HA) have been shown to form a pro-regenerative matrix in newt heart (48). Thus, a comprehensive study of the zebrafish heart ECM composition and characteristics may reveal other key points in the heart regeneration process. In the present study, we have developed a protocol to decellularize adult zebrafish hearts, and applied it to non-injured and regenerating hearts at 7, 14 and 30 days post-amputation (dpa). We have then characterized the composition of the zebrafish heart ECM, as well as the major changes in ECM composition that take place during heart regeneration. Finally, we have used atomic force microscopy (AFM) to analyze the effect that changes in ECM composition have on matrix stiffness. Our studies identify important changes in ECM protein composition and mechanical properties during zebrafish heart regeneration.

## Results

### Enrichment of zebrafish heart ECM proteins

To study the ECM protein composition of zebrafish ventricles, we developed a decellularization protocol consisting on detergent treatment with 0,5% sodium dodecyl sulfate (SDS), followed by 1% Triton-X 100 to remove SDS prior to processing for analysis (Fig. 1A). This treatment produced translucent decellularized scaffolds that structurally resembled the zebrafish ventricle (Fig. 1B). Following decellularization, DAPI staining and other histological stainings were used to assess the extent of cell removal and the integrity of the ECM. No DAPI staining was seen after treatment (Fig. 1C), and spectrophotometric analysis showed an ∼80% reduction in DNA content after decellularization of zebrafish hearts (Fig. 1D). Moreover, hematoxylin and eosin, and Masson trichrome stainings revealed no conspicuous damage to the ECM, while further confirming the absence of nuclei in decellularized samples (Fig. 1E). To analyze whether the decellularization process altered the relative abundance of ECM proteins, we characterized the proteome of zebrafish hearts before (native), after SDS treatment, and at the end of decellularization by liquid chromatography-mass spectrometry (LC-MS). A total of 447 unique proteins were detected in native hearts, of which only 15 (3.4%) corresponded to extracellular proteins as annotated by Gene Ontology and manual curation (see Experimental Procedures for details). The numbers of extracellular proteins detected after SDS treatment and at the end of the decellularization protocol increased to 35 and 36, respectively, while those of intracellular proteins decreased from 432 in native hearts to 352 after SDS treatment, and 341 in fully decellularized hearts (Supplementary Table 1). As expected, decellularization not only decreased the number of intracellular proteins detected, but also their abundance as quantified by peptide peak intensity, whereas the opposite trend was observed for extracellular proteins (Fig. 1F). Importantly, all extracellular proteins identified in native hearts were also detected in samples after SDS treatment, and after complete decellularization, demonstrating that the protocol used for decellularizing zebrafish hearts did not introduce bias in our analyses due to preferential removal of specific ECM components. The end result of the decellularization protocol was, therefore, an enrichment of ECM proteins at the expense of cytoplasmic and nuclear proteins (Fig. 1G). The results of these analyses validate the applicability of our zebrafish heart decellularization protocol to enrich for ECM proteins, while not altering the relative abundance of ECM protein components.

**Table 1.**
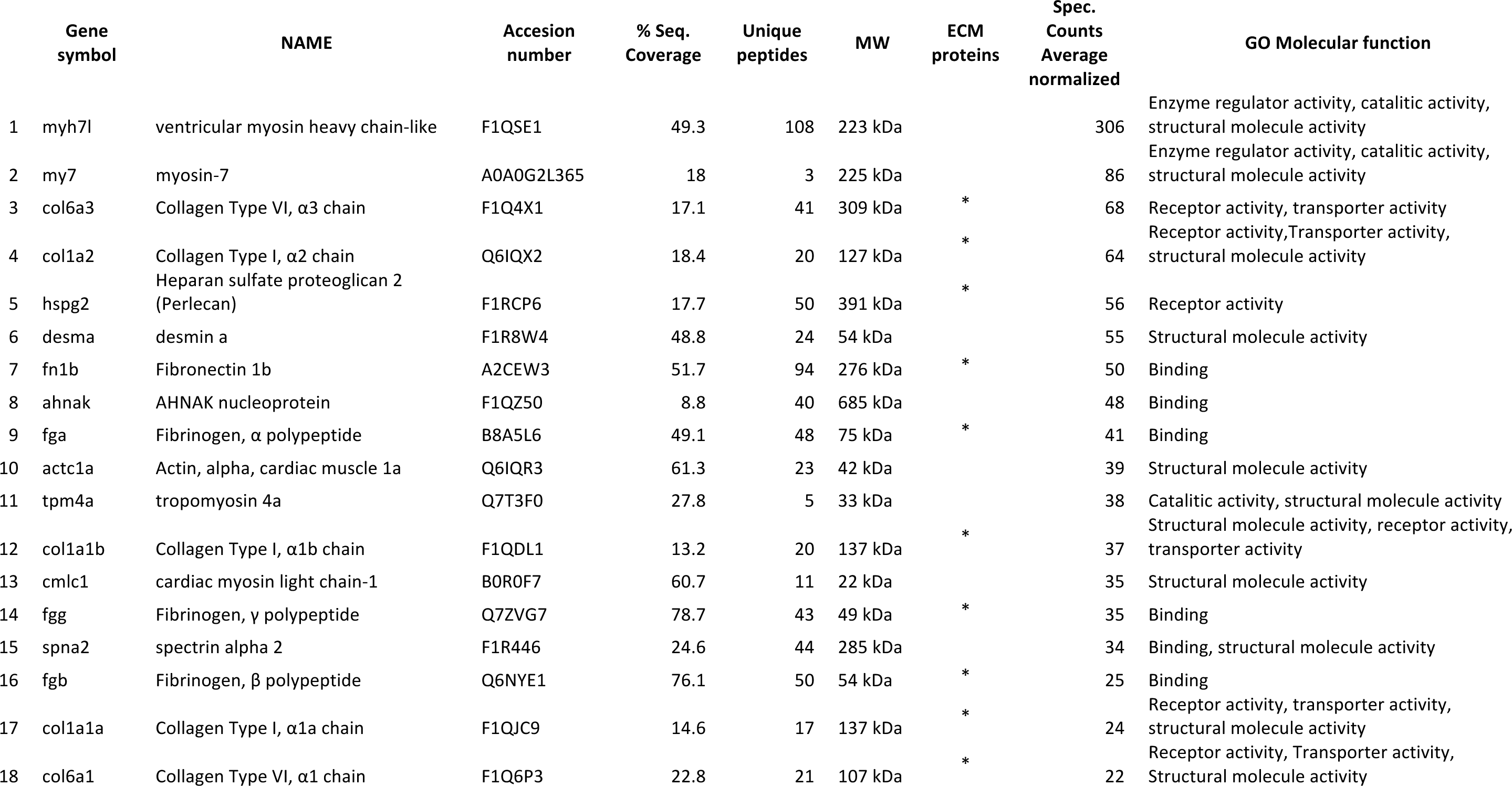
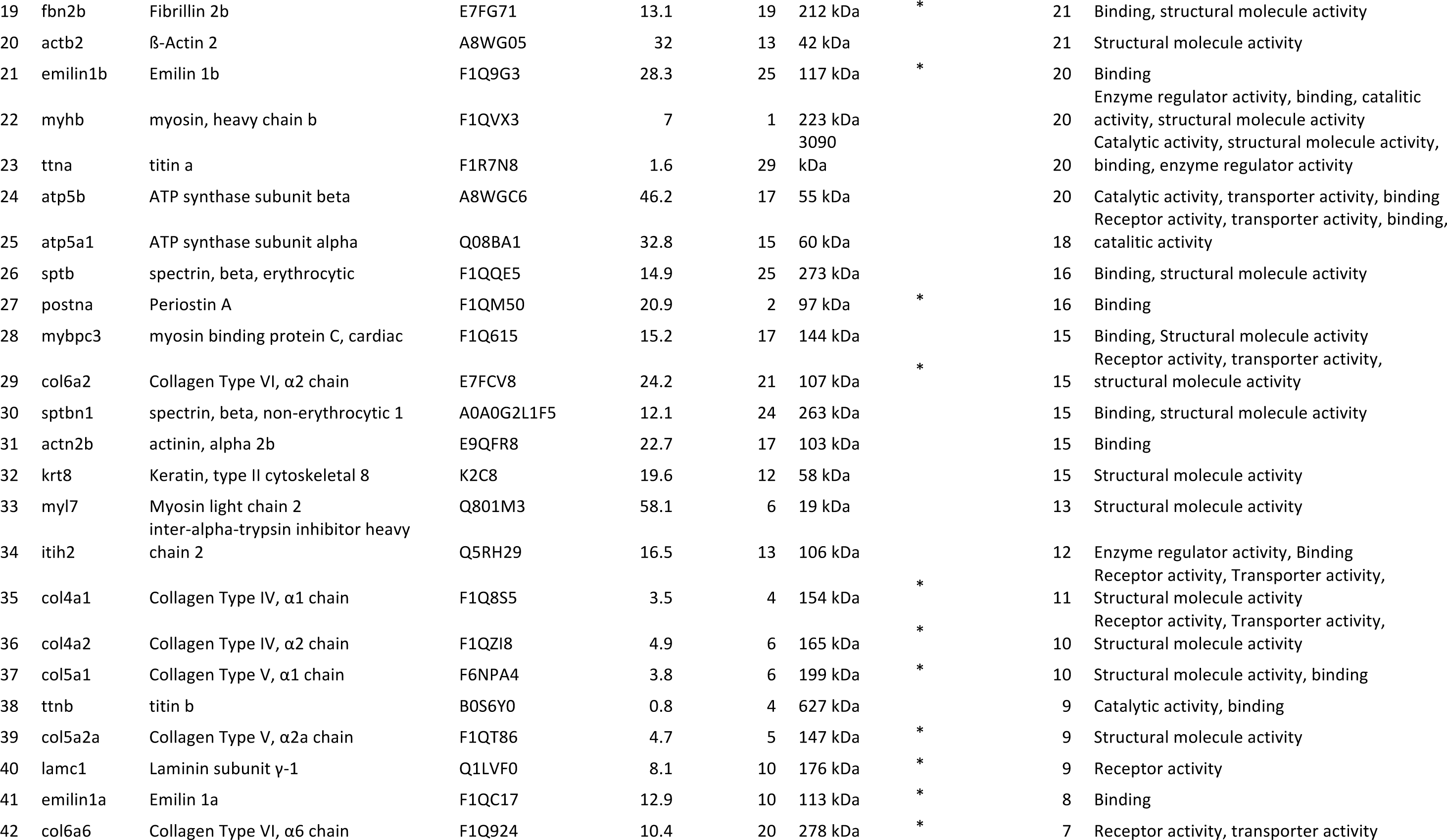
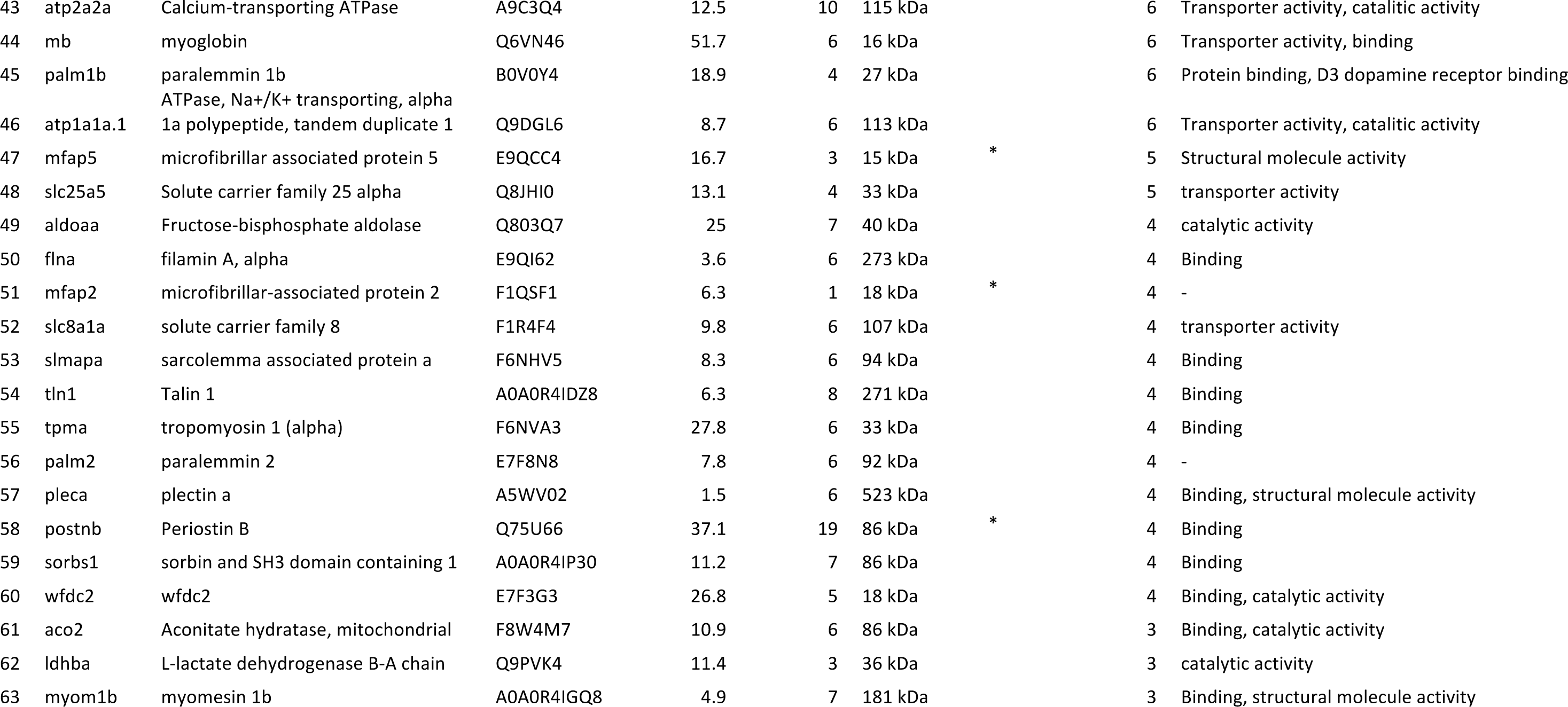
Proteome of decellularized zebrafish ventricles. List of proteins identified in the proteomic analysis that were represented over 3 spectral counts. Spectral counts were normalized by the total spectral counts in the proteomic analysis. MW, molecular weight. GO, gene ontology.

**Figure 1.**
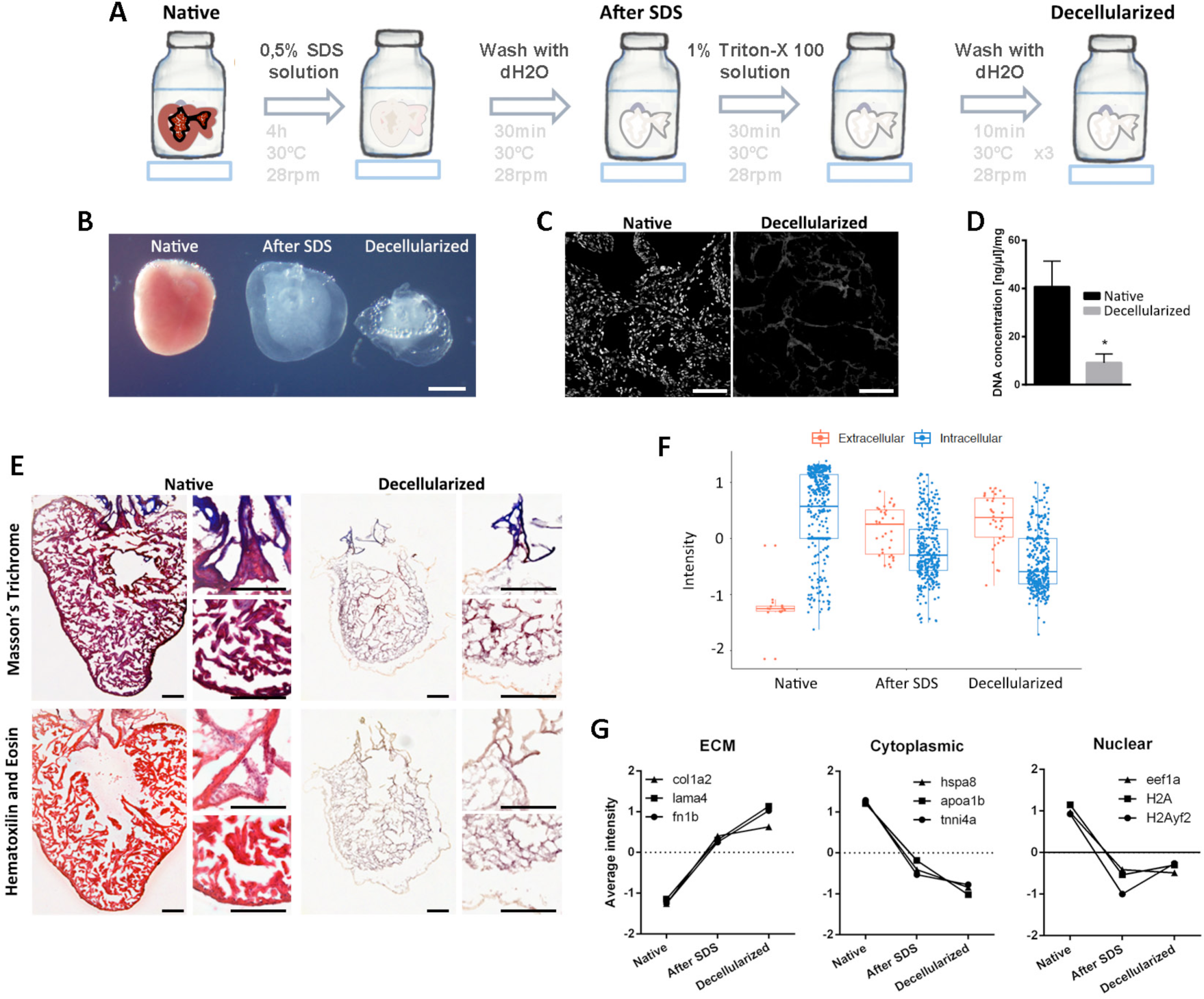
Decelularization protocol for zebrafish hearts. (A) Representation of the protocol used to decellularize zebrafish ventricles. (B) Images of the ventricles at different points of the decellularization protocol. From left to right, native ventricle, after 0.5% SDS, and at the end of the decellularization ptotocol (scale bar 500µm). (C-E) The efficiency of the decellularization process was characterized by DAPI staining (scale bar 50µm) (C), spectrophotometric quantification of DNA (N=3) (D), and Masson’s trichrome (E, upper images) and Hematoxilin and Eosin (E, lower images) staining (scale bar 100µm). (F) Box plot representing an overall distribution of the average intensities of each protein in each sample group. ECM proteins and intracellular proteins have been independently plotted to better visualize the effect of decellularization in each class of proteins. (G) The profile of specific ECM, cytoplasmic and nuclear proteins is plotted to indicate the loss of intracellular proteins while the maintenance and enrichment of ECM proteins. Statistical significance of DNA content was analyzed with unpaired Student’s *t* test. *, p<0.05.

### Profile of ECM proteins in adult zebrafish hearts

Proteomics analysis of the decellularized sham zebrafish hearts by LC-MS allowed us to identify a total of 63 proteins with ≥3 spectral counts, of which 24 were ECM proteins (Table 1). Collagens, fibronectin 1b and fibrinogens were the most abundant proteins in the decellularized zebrafish ventricle proteome. Binding, structural molecule activity, receptor activity, and catalytic activity were the most represented molecular function Gene-Ontology (GO) terms (GO:0005488, GO:0005198, GO:0004872, and GO:0003824, respectively) for these identified ECM proteins. A search for GO enriched terms using Enrichr analysis revealed that the 10 most enriched biological processes in our control proteome (GO:0030199, GO:1905590, GO:0071711, GO:0030198, GO:0098784, GO:0010215, GO:0061148, GO:1901148, GO:0021820 and GO:0022617), were all ECM related (Fig. 2A).

**Figure 2.**
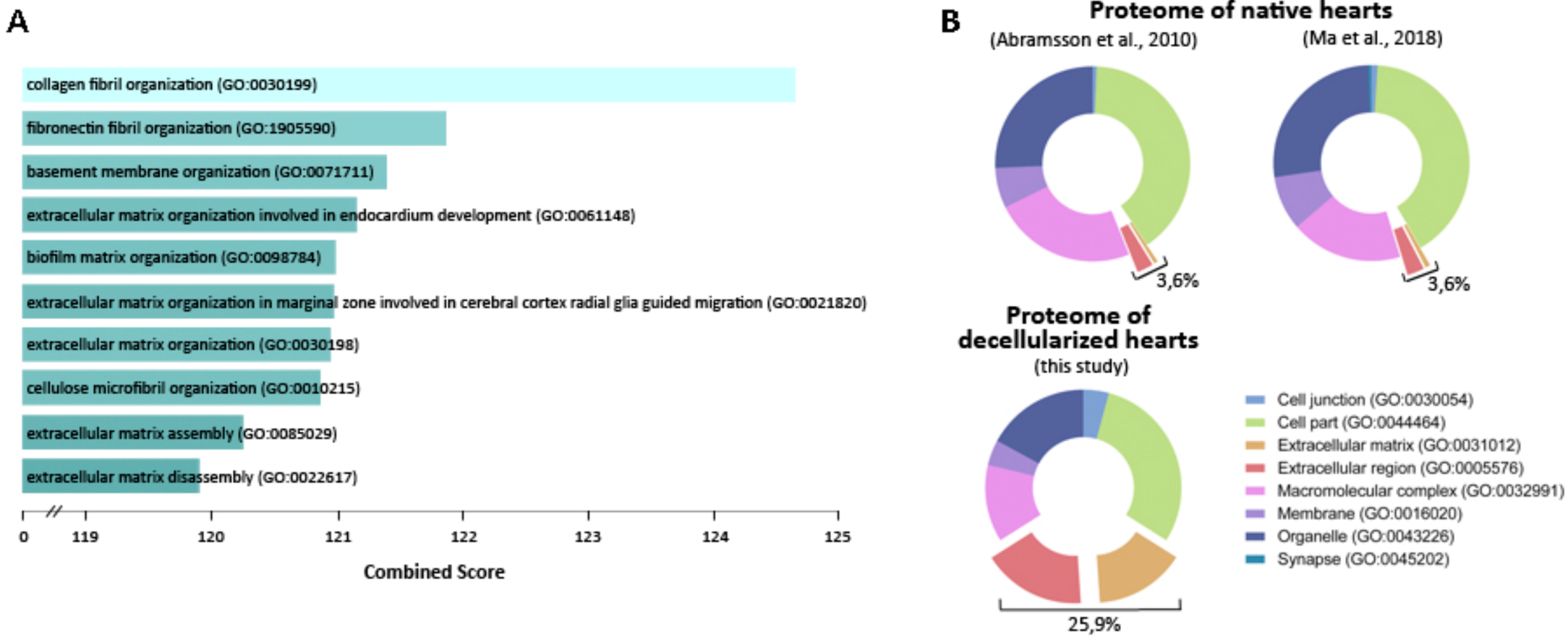
Decellularized zebrafish ventricles are enriched in ECM proteins. (A) Representation of the 10 most enriched Biological Process GO terms in our proteome of control hearts. Combined score is taken from Enrich program, which takes the log of the p-value from the Fisher exact test and multiplies it by the z-score of the deviation from the expected rank. (B) Representation of the Cell Component Gene Ontology (GO) of the published zebrafish heart proteome of Abramsson et al. (2010) (49), the published sham heart proteome of Ma et al. (2018) (50) and our proteomic analysis of decellularized zebrafish ventricles. Panther assigned GO terms to 138, 1902, and 60 proteins from the proteomes described in Abramsson et al. (2010) (49), Ma et al. (2018) (50), and the one described here, respectively.

We next compared the GO cellular component terms represented in our ECM-enriched zebrafish ventricle proteome with those of whole cardiac zebrafish proteomes previously reported by Abramsson et al. (49) and Ma et al. (50) Analysis by Panther revealed an enrichment of ECM and extracellular region terms in our decellularized samples compared to whole heart proteomes (Fig. 2B). These results further confirmed that ventricle decellularization allows a better detection of changes in ECM protein composition.

### Changes in ECM protein composition during zebrafish heart regeneration

To determine whether ECM protein composition changes during zebrafish heart regeneration, we also analyzed with LC-MS decellularized ventricle samples of zebrafish hearts at 7, 14, and 30 days post-amputation (dpa). From the 274 proteins detected among all samples (Supplementary Table 2), 96 were represented by an overall across sample averages of >5 spectral counts (Supplementary Tables 3 and 4). From these 96 proteins, 29 corresponded to extracellular proteins and 67 to intracellular proteins. Of note, 23 out of the 50 most abundant proteins found in these analyses were ECM proteins (Supplementary Tables 3 and 4). Heatmap representation of protein profiles showed large differences between control and 7-dpa samples. However, profile changes among other regeneration time points were less evident (Fig. 3A). Overall, hierarchical clustering analysis grouped sample replicates together for all time points with good Approximately Unbiased p-values (AU) indexes, except for those of the 14-dpa condition. Control and 30-dpa samples clustered together, and away from 7-dpa samples, while the 14-dpa samples clustered apart from each other (Fig. 3B). ANOVA analysis revealed that 17 out of the overall 96 proteins detected showed statistically significant changes during regeneration, of which 9 were ECM proteins (Supplementary Table 5). Among the ECM proteins whose levels changed significantly during regeneration, fibrinogen a, b, and g, as well as fibronectin 1b and periostin b, showed a peak at 7 dpa. In contrast, the levels of collagens and fibrillin 2b showed a statistically significant decrease during regeneration, which was more evident at 7 dpa.

**Figure 3.**
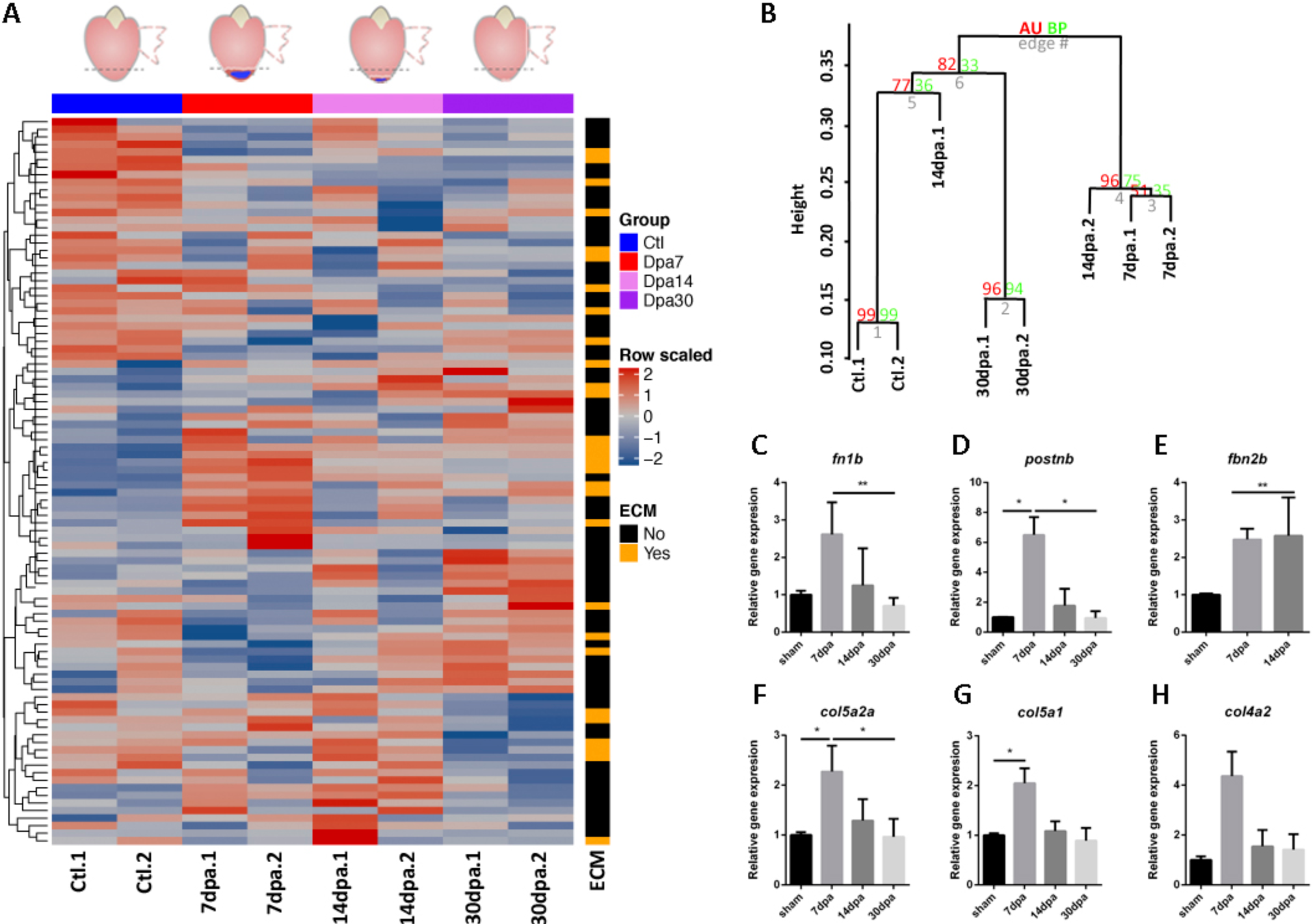
Changes in ECM protein composition during heart regeneration. (A) Heat-map for the 96 proteins detected across samples in zebrafish control decellularized hearts (Ctl) and at different time points of regeneration. The time points analyzed were 7 days post-amputation (dpa), 14 dpa, and 30 dpa. Red indicates increased protein expression and blue indicates reduced protein expression. The ECM column indicates the ECM proteins in orange. Data are row scaled. (B) Hierarchical clustering with bootstrap analysis of all the samples. AU, Approximately Unbiased p-value; BP, Bootstrap Probability value. (C-H) Gene expression assessment by real time qPCR of the ECM proteins significantly changing in the proteomic analysis during the regeneration process. *fn1b*, fibronectin 1b; *postnb*, periostin b; *col5a1*, collagen type 5 α1 chain; *col4a2*, collagen type 4 α2 chain; *col5a2a*, collagen type 5 α2a chain; *fbn2b*, fibrillin 2b. Significance was analysed with Kruskal-Wallis followed by a Dunn’s multiple comparisons test. *, p<0.05; **,p<0.01.

To ascertain if changes in protein abundance were the result of differential gene expression, we measured by quantitative reverse transcription polymerase chain reaction (qRT-PCR) the expression level of 6 genes encoding ECM proteins that change in abundance during regeneration. We found positive correlation between mRNA and protein levels in the case of fibronectin/*fn1b* and periostin b/*postnb*, in which increased transcription levels were also found peaking at 7 dpa (Fig. 3C-D). However, expression levels of *col4a2, col5a1, col5a2a*, and *fbn2b* were all found to be upregulated at 7dpa (Fig. 3E-H), in contrast with the decreased protein levels found in regenerating heart ECM (Supplementary Table 5). This suggests that changes in fn1b and postnb protein levels during regeneration are regulated transcriptionally, whereas the regulation of *col4a2, col5a1, col5a2a*, and *fbn2b* would be post-transcriptional.

### Changes in ECM biomechanical properties during zebrafish heart regeneration

Changes in ECM protein composition can result in modifications in the biomechanical properties of the matrix. Atomic Force Microscopy (AFM) allowed us to measure and compare the stiffness of the ECM during heart regeneration (Fig. 4). For this purpose, 25-µm thick slices of control or regenerating zebrafish hearts were decellularized and the ECM regions of interest identified by phase contrast microscopy (Fig. 4A-C). The Young’s modulus of control zebrafish ventricle ECM was calculated at 8.1 ± 1.7 kPa. We then compared these values with those of regenerating hearts at 7 and 14 dpa (Fig. 4D). We chose these time points because our proteomic analysis had revealed the major ECM changes at 7 dpa (Fig. 3A and B, and Supplementary Tables 3 and 4). AFM measures of the non-injured myocardium and the regenerating area revealed an overall significant decrease of the ventricular ECM stiffness at 7 dpa (myocardium 2.8 ± 0.5 kPa and wound 3.7 ± 1.0 kPa). The stiffness of decellularized ventricles returned back to control values by 14 dpa (myocardium 14.4 ± 4.8 kPa and wound 11.9 ± 4.0 kPa). No significant differences were detected between the non-injured myocardium and the regenerating area in any time points analyzed, suggesting that the ECM changes observed take place at the organ level.

## Discussion

Extracellular matrix (ECM) remodeling is a critical step in development, wound healing and regeneration (31, 38, 40, 41, 51). The present study characterized the ECM composition and its changes during zebrafish regeneration. We have developed a decellularization protocol for zebrafish ventricles that results in ECM enrichment, as well as facilitating the analysis of the proteomic profile of the zebrafish ventricle ECM. Moreover, we have analyzed the ECM changes during heart regeneration and assessed the stiffness of the ECM at different time points of this process. Altogether, the results from our studies should help better understanding the role of the ECM in zebrafish heart regeneration.

**Figure 4.**
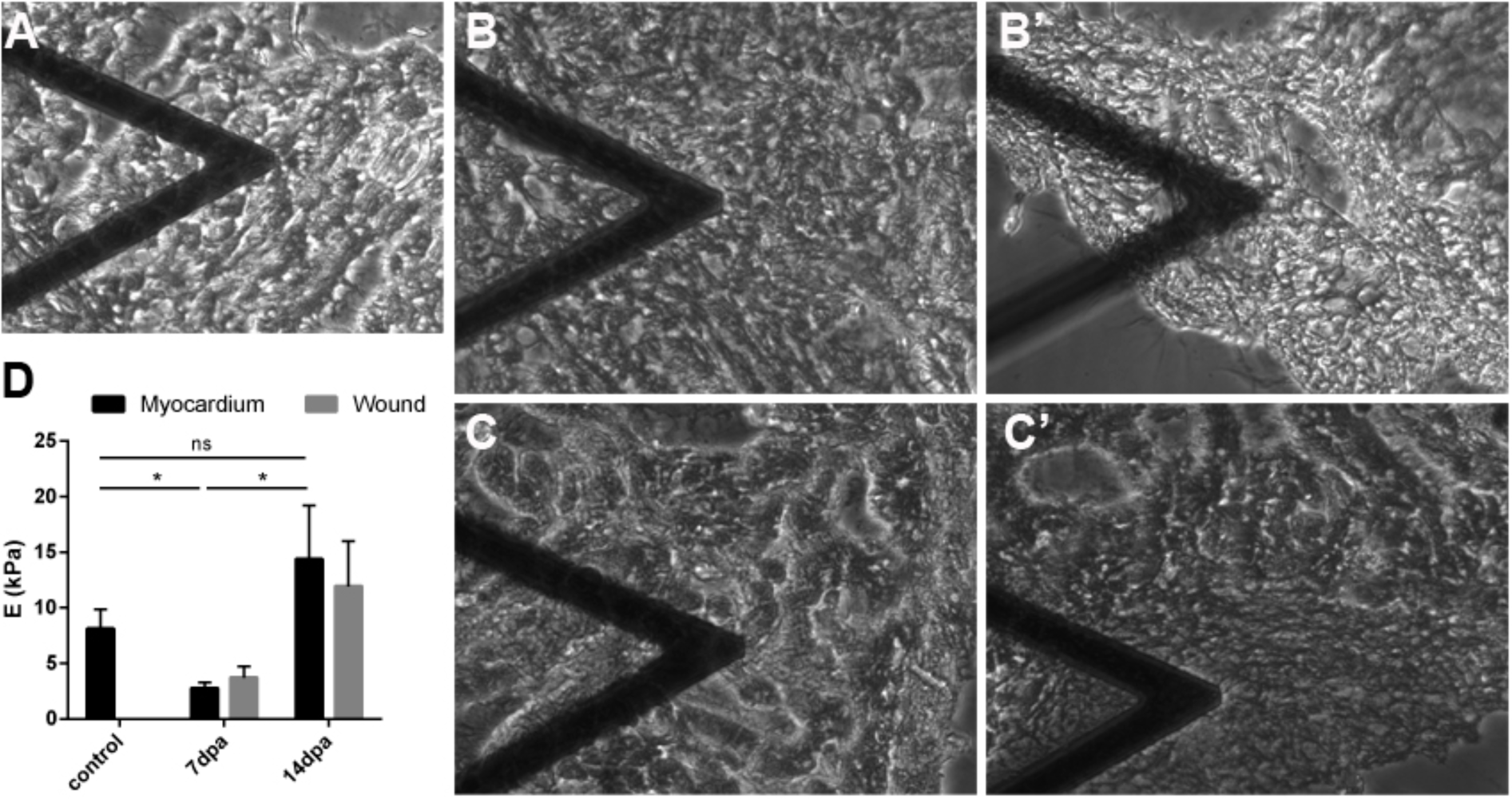
Stiffness of the extracellular matrix of regenerating hearts. (A-C) Bright field images of decellularized zebrafish hearts being analyzed by AFM at the myocardium away from the injury area. (B’-C’) Bright field images of decellularized zebrafish hearts being analyzed by AFM at the regenerating area. (A) Control decellularized heart, (B,B’) 7 dpa decellularized hearts, (C,C’) 14 dpa decellularized heart. The triangular shape in A-C and B’-C’ is the AFM cantilever. (E) Young’s modulus of the heart ECM of non-injured, 7 dpa, and 14dpa hearts (N=5 each). dpa, Days post-amputation. Statistically significance was assessed with Man-Whitney test. *, p<0.05

The ECM composition has not been fully studied and described in the specific contexts of the zebrafish heart and cardiac regeneration. Few are the studies done to analyze the ECM in the zebrafish heart. The proteome of different zebrafish organs, including the heart, were analyzed by Abramsson and colleagues and only 4 collagen proteins were detected in the heart proteome (49). In the present study, we have described the presence of 7 additional collagens (col1a2, col1a1b, col6a2, col4a1, col4a2, col5a2a, and col6a6) in control samples, corroborating that they are the main structural element of the zebrafish heart ECM. Collagens provide tensile strength, regulate cell adhesion, support chemotaxis and migration, and direct tissue development (52). Recently Chen and colleagues analyzed the decellularized ECM of zebrafish heart using a mechanical decellularizing approach (53). They qualitatively described the presence of 4 ECM proteins, of which we detect 3 as well as 21 new ones. Thus our decellularization process followed by LC/MS analysis provided a more comprehensive, as well as quantitative, method to define the ventricular ECM of the zebrafish.

In terms of the cardiac regeneration process, there was no study to our knowledge that analyzed the importance of the ECM itself in the zebrafish cardiac regeneration model. Transcriptomic and proteomic approaches have sought to profile the whole gene expression and protein changes during zebrafish heart regeneration. The former identified gene expression changes during regeneration process and described an increase of transcripts related with secreted molecules, catepsins and metalloproteinases, and wound response/inflammatory factors (45, 46). Also, transcripts coding for ECM and adhesion molecules were detected to be commonly and differentially expressed when comparing the transcriptomes of zebrafish regenerating hearts and fins (46). On the other hand, proteomic studies have been mainly done on native cardiac samples without decellularization prior to protein detection, where ECM proteins could be masked by the large amount of intracellular proteins (31, 32, 50, 53). Moreover, these previous proteomic studies have focused on early regenerating time points and do not provide an overview of the proteomic changes during the entire regenerating process (32, 53).

It is also worth noting that, in our studies, ECM proteins were overrepresented among those that showed significant changes in abundance during heart regeneration. Thus, enrichment analysis using Enrichr tool on all proteins that showed significant changes during regeneration identified Biological Process GO terms related with ECM as the most represented (Supplementary Fig. 1A). In contrast, a similar analysis of published proteomic data (50) only identified enrichment of the ECM-related GO terms fibrinolysis (GO:0042730) and plasminogen activation (GO:0031639), and not among the most enriched in that dataset. A summary of the 10 most-enriched Biological Process GO terms during zebrafish heart regeneration identified in our studies and in those of Ma and colleagues (50) is presented in Supplementary Fig. 1B.

Both transcriptomic and proteomic studies have identified some ECM proteins important during regeneration. Wang and colleagues described the importance of fibronectin during regeneration after ventricular resection, suggesting a positive effect on cardiomyocyte migration (31). Also, tenascin C has been found to be expressed at the border zone and suggested to mediate loosening of cardiomyocyte attachment to the substrate, thereby facilitating cardiomyocyte migration (21). Another proteomic study identified the hyaluronic acid receptor (Hmmr) to be important for the epicardium EMT migration towards the injury (32). All these suggest that tissue remodeling and ECM dynamics are important factors during zebrafish heart regeneration.

The fact that our analyses did not detect changes in some ECM proteins found in previous studies, such as tnc, may be due to technical limitations. In LC-MS-based analyses, signals of abundant proteins such as collagens and structural proteins can mask the signals of low-abundance proteins. Also, new proteomic analytical tools have been developed, but we strongly believe that proteomic analysis of ECM enriched samples is a powerful approach to study the ECM components in heart regeneration. In this study we have identified the main changes in ECM protein composition during zebrafish heart regeneration. Decreased amounts of collagen IV, collagen V, collagen VI, and fibrillin 2b, as well as increased amounts of fibrinogens, fibronectin 1b and periostin b comprise the initial regenerating ECM. We have been also able to detect proteins that were not previously detected in any transcriptomic or proteomic study, such as col4a2 and fbn2b. Two groups of proteins appear to be regulated differently. In the first group, positive correlation between protein abundance and gene expression in the case of fn1b and postnB, indicates a transcriptional regulation of these genes during heart regeneration (37, 46). In the second group, decreased levels of col4a2, col5a1, col5a2a and fbn2b proteins inversely correlate with gene expression, which we interpret as a regulatory feedback mechanism in an attempt to recover the protein levels.

Stiffness is itself a mechanical property of the ECM. It is known to be involved in cell processes such as the regulation of cell proliferacion (43, 54), dedifferentiation (43), migration (through durotaxis) (55) and stem cell differentiation (56). We analyzed the stiffness of the regenerating zebrafish ventricle and found a significant decrease in ECM stiffness at 7 dpa. It is known that an increase in collagen concentration or cross-linking is associated with stiffer myocardium (57), whereas an extensive degradation of myocardial collagen is associated with a decrease in ventricular stiffness (57, 58). Thus, this result correlates well with the decrease in abundance of several collagens identified in our proteomic analyses, although further research will be necessary to ascertain if these two findings are, indeed, causally related. An unexpected finding of our mechanical characterization of the regenerating heart ECM was that we did not detect any conspicuous differences in stiffness between the regenerating area and the non-injured myocardium located far away from the lesion. This suggests that changes in ECM composition result in organ-wide effects at the biomechanical level. Interestingly, recent data from our laboratories have identified a low ECM stiffness as a permissive factor regulating the heart regeneration ability of neonatal mice (59). The exact mechanism(s) by which ECM stiffness regulates heart regeneration competence in adult zebrafish and/or neonatal mice require further investigation.

Changes in ECM composition and stiffness are likely to instruct specific cell behaviors, and may also trigger further changes in the properties of the ECM itself. The ability of the zebrafish cardiac ECM to induce mammal heart regeneration has been very recently assessed by Chen and colleagues (53). They observed that the zebrafish cardiac ECM exhibited pro-proliferative and chemotactic effects *in vitro* and contributed to a higher cardiac contractile function in mouse. In general, fibrillins are known to play important roles in TGFβ signaling by controlling the amounts of cytokines and in endocardium morphogenesis (60, 61). Signaling by the TGFβ/Activin pathway is known to promote cardiomyocyte proliferation and deposition of ECM proteins (62). Moreover, fibronectin 1b promotes cardiomyocyte migration during zebrafish cardiac regeneration (31). Periostin stimulates healing after myocardial infarction in mice through induction of cardiomyocyte proliferation, and it is known to be responsible of collagen cross-linking (covalent linkage of collagen fibers) (63, 64). Collagen cross-linking, in turn, increases at the same time the accumulation of collagen, matrix rigidity and resistance to degradation (65, 66). Further studies to clarify in detail the function of each ECM protein during cardiac regeneration may lead to a better understanding of this process, and to the development of new avenues for therapeutic intervention to promote regeneration of the mammalian heart.

## Methods

### Animal maintenance and surgical procedure

Wild-type zebrafish of the AB strain were maintained according to Standard protocols (67). The ventricular amputations were done as previously described by Raya et. al, 2003). Animal procedures were performed under the approbation of the Ethics Committee on Experimental Animals of the PRBB (CEEA-PRBB).

### Ventricle decellularization

Animals were sacrificed and hearts were extracted after an intra-abdominal injection of 20µl of heparin (1,000U/ml, Sigma). Only the ventricles were decellularized by immersion in a 0.5% SDS solution for 4h. Then ventricles were washed with distilled H_2_O for 30min. Then immersed into a 1% Triton-X solution for 30min and finally washed three times with distilled H_2_O for 10min each wash. All solutions were previously filtered with a 0.2µm filter, and all incubations steps were done in a horizontal shaking plate at 30°C and 28 rpm.

### Tissue processing and staining

Zebrafish hearts were extracted as previously mentioned and fixed overnight with 4% paraformaldehyde. Then the hearts were incubated overnight at 4°C with 30% sucrose and frozen in OCT (Tissue-Tek) for cryosectioning. The decellularized ventricles were embedded in OCT, snap frozen with isopentane and fixed after sectioning incubating them 10min in 4% paraformaldehyde. 10µm thick slices of the samples were counterstained with DAPI (1:10,000) for 4min and stained with Hematoxylin and Eosin, and Masson’s Thrichrome stainings.

### Genomic DNA extraction and quantification

Genomic DNA was extracted from non-decellularized and decellularized zebrafish ventricles. Samples were homogenized by adding 200µl of PBS, 20µl Proteinase K (Qiagen) and 4ul of RNAseA (Qiagen) and vortexed. Tissue lysis was done adding 200µl of AL Lysis buffer (Qiagen). DNA was purified by chloroform and precipitated using isopropanol. Finally the pellets were dried and 20µl of TE buffer (Quiagen) was added. DNA concentration was measured with a spectrophotometer (NanoDrop® ND-100, Thermo Fisher Scientific).

### Liquid chromatography-Mass spectrometric analysis

Two different proteomic analyses were performed: one to assess the decellularization protocol, and another one to assess the regeneration process. For the first one, native zebrafish ventricles, half-decellularized ventricles, and fully decellularized ventricles were analyzed. For the second one, decellularized zebrafish ventricles at 7days post-amputation (dpa), 14dpa and 30dpa, were analysed compared to sham operated fishes. Proteins of each sample were solubilized by mixing with 50uL of 1% SDS, 100mM Tris-HCl pH 7.6, 100mM DTT, 10min sonication and boiling for 3min. Protein extracts were clarified by centrifugation at 16,000xg for 5min, and quantified using the RcDc kit (BioRad). In the first proteomic analysis, 12 μg of protein of each sample were digested with LysC and trypsin using a Filter-Aided Sample Preparation (FASP) protocol and further analyzed by mass spectrometry. The LC separation was conducted on an Easy-nLC 1000 (Thermo) using 0.1% formic acid as Solvent A and acetonitrile with 0.1% formic acid as B. Each run, was carried out at 250 nL/min with a gradient of 95% of solvent A to 65% A in 180 min. Blank samples with solvent A injections were run in between each sample. Sample was concentrated in an Acclaim PepMap 100 trap column (Thermo), and fractionated in a Nikkyo Technos Co., 75 um ID, 3 A pore size, 12.5 cm in length with built in emitter column, coupled to a Nanospray Flex (Thermo) ESI source. Shotgun LC-MS/MS analysis was performed online with an electrospray voltage of 1.9 kV using a Q Exactive HF mass spectrometer (Thermo) with HCD fragmentation using top 15 precursor with charge 2 to 5 for data-dependent acquisition (DDA). MS1 spectra were acquired in the mass range 390-1700 m/z at a resolution of 60,000 at m/z 400 with a target value of 3×10^6^ ions and maximum fill time of 20 ms. MS2 spectra were collected with a target ion value of 2×10^5^ and maximum 100 ms fill time using a normalized collision energy of 27. Dynamic precursor exclusion was set at 15s. The raw files were processed with the MaxQuant software (version 1.6.2.6a) using the built-in Andromeda Search Engine. The *Danio rerio* TrEMBL database downloaded from www.uniprot.org (Oct, 8th 2018) (62,078 entries) was used to search for peptides. MS/MS spectra were searched with a first search precursor mass tolerance of 20 ppm. Then, the peptide masses were corrected and a second search was performed at 4.5 ppm of mass tolerance. The fragment tolerance was set to 0.5Da, the enzyme was trypsin and a maximum of 2 missed cleavages were allowed. The cysteine carbamidomethylation was set as fixed modification and methionine oxidation as well as protein N-terminal acetylation as variable modifications. To improve the identifications the “match between runs” was enabled among the replicates of every experimental condition.

In the second proteomic analysis, the protein amount recovered was around 5 microgram for the 0.5% SDS treated sample. The buffer was changed to 2M Urea 50mM Ammonium Bicarbonate using a 5kD Amicon Ultrafiltration device, and the samples were digested with trypsin. Each sample was the analyzed by LC-MS in duplicate. 500ng of each sample was analyzed on a Maxis Impact high-resolution Q-TOF spectrometer (Bruker, Bremen), coupled to a nano-HPLC system (Proxeon, Denmark). The samples, evaporated and dissolved in 5% acetonitrile, 0.1% formic acid in water, were first concentrated on a 100mm ID, 2cm Proxeon nanotrapping column and then loaded onto a 75mm ID, 25cm Acclaim PepMap nanoseparation column (Dionex). Chromatography was run using a 0.1% formic acid - acetonitrile gradient (5-35% in 120min; flow rate 300nL/min). The column was coupled to the mass spectrometer inlet through a Captive Spray (Bruker) ionization source. MS acquisition was set to cycles of MS (2Hz), followed by Intensity Dependent MS/MS (2-20Hz) of the 20 most intense precursor ions with an intensity threshold for fragmentation of 2500 counts, and using a dynamic exclusion time of 0.32min. All spectra were acquired on the range 100-2200Da. LC-MSMS data was analyzed using the Data Analysis 4.0 software (Bruker). Proteins were identified using Mascot (ver. 2.5; Matrix Science, London UK) to search against the *Danio rerio* proteins in the SwissProt 20160108 database (43,095 sequences). MS/MS spectra were searched with a precursor mass tolerance of 10ppm, fragment tolerance of 0.05Da, trypsin specificity with a maximum of 2 missed cleavages, cysteine carbamidomethylation set as fixed modification and methionine oxidation as variable modification.

Both mass spectrometry proteomics datasets, for the decellularization protocol and for the regeneration proteome, have been deposited to the ProteomeXchange Consortium (http://proteomecentral.proteomexchange.org) via the PRIDE partner repository (68) with the dataset identifiers <PXD011627> and <PXD010092>, respectively.

### Criteria for protein identification

For the decellularization protocol proteomic analysis, MaxQuant software (version 1.6.2.6a) was used to validate the peptides and proteins identifications. The final list of peptides was obtained after applying a 5% False Discovery Rate (FDR). For proteins, only the proteins with at least 1 assigned peptide after applying a 5% FDR were considered. The non-unique peptides were assigned to the corresponding protein group according to the Razor peptides rule implemented in the software (principle of parsimony). Finally, the identified peptides and proteins were filtered to remove the peptides/proteins tagged as “Reverse” (significantly identified in the reverse database) and “potential contaminant” (items identified as contaminants in the “contaminants.fasta” file) as well as the proteins “Only identified by site” (proteins identified only with modified peptides). The lists can be found in the supplementary material uploaded to the PRIDE repository, with project accession code <PXD011627>.

For the regeneration process proteomic analysis, Scaffold (version Scaffold_4.0.5, Proteome Software Inc., Portland, OR) was used to validate MS/MS based peptide and protein identifications. Peptide identifications were accepted if they could be established at greater than 99% probability by the Peptide Prophet algorithm (69) Protein identifications were accepted if they could be established at greater than 98% probability to achieve an FDR less than 1% and contained at least 1 identified peptide. Protein probabilities were assigned by the Protein Prophet algorithm (70). Protein isoforms and members of a protein family would be identified separately only if peptides that enable differentiation of isoforms had been identified based on generated MS/MS data. Otherwise, Scaffold would group all isoforms under the same gene name. Different proteins that contained similar peptides and which were not distinguishable based on MS/MS data alone were grouped to satisfy the principles of parsimony. The lists of identified peptides and proteins can be found in Supplementary Tables 6 and 7, respectively.

### Label-free Protein Quantification

For the decellularization protocol proteomic analysis, the proteins were quantified with the help of the label-free algorithm (LFQ) implemented in the MaxQuant software using the unique and razor peptides. The minimum number of peptides to be available in all the pair-wise comparisons was set to 2 and the “stabilize large LFQ ratios” option enabled.

For the proteomic analysis of heart regeneration, relative label-free protein quantification analysis was performed on the different samples analyzed using spectral counting. The “Quantitative Value-Total Normalized Spectra” function of Scaffold software was used for quantitative comparison. This function provides the total number of spectra that matched to a protein identified in each sample, after normalizing the values for each sample by a factor calculated so that the total number of normalized spectral counts is identical for all samples, thus correcting for differences in sample load. Only those proteins for which the total sum of spectral counts for the eight runs was greater than 5 were considered for quantitative comparison.

### Analysis and validation of proteomic data on regeneration

In order to annotate proteins associated with Extracellular Matrix we downloaded the GO annotations of zebrafish, human, and mouse organisms from the repository of Gene Ontology Consortium (71, 72). The resulting list of proteins falling into the GO Term GO:0031012 (i.e. Extracellular Matrix) was supervised and curated by hand. To determine an enrichment in ECM proteins in or decellularized samples, the proteomics data obtained was compared to the zebrafish proteomic data already published (49) with Panther (pantherdb.org) for ECM protein enrichment. GO enrichment analysis were done with Enrichr tool (version August 24^th^, 2017). To stabilize the variance, Variance Stabilization Normalization (VSN) was applied to spectral counts (73). A hierarchical clustering analysis via 1,000 bootstrap resampling replications was performed, using the R (version 3.2.3) statistical language package pvclust (74), to construct a dendrogram with correlation distance and average method. Normalized data values were z-scaled by rows in order to build the heatmap. Validation of the proteomic data was performed by qRT-PCR analyzing 3 biological replicates comprised of 5 ventricles each for each time point (7dpa, 14dpa, 30dpa and sham). RNA was extracted with TriReagent (Molecular Research Center, Inc.) and chloroform, and cDNA was synthesized using the Transcriptor First Strand cDNA Synthesis Kit (Roche).

### Atomic force microscopy (AFM) for the measurement of the extracellular matrix

Zebrafish hearts were extracted, included in Optimal Cutting Temperature coumpound (OCT) and freezed at −80 °C. Then slices of 25µm were cut with a cryostat (HM 560, CryoStar Thermo Scientific) and placed on top a positively charged glass slides and stored at −20 °C. Before measurements, slices were washed with PBS 1x and decellularized in order to remove all cellular components and preserve the ECM. A detergent-based protocol was used to decellularize the slices. Briefly, slices were immersed in sodium-dodecyl disulfate (SDS) 0.1% during 5 minutes, followed by 10 min of Triton X-100 1% and finally washed during 20 min with NaCl 0.9% solution. The slices were kept in constant moderate agitation.

Immediately after decellularization process, the samples were measured by AFM. The perimeter of glass slides was outlined with a water repellent marker (Super PAP PEN, Invitrogen), keeping 1 ml of PBS 1x pooled over the slices. Then, slides were placed on the sample holder of a custom-built AFM coupled to an inverted optical microscope (TE 2000, Nikon). The Young’s modulus (E) of the ECM was measured using V-shaped Au-coated silicon nitride cantilever (nominal spring constant of k = 0.06 N/m) with a spherical polystyrene bead of 2.25μm radius glued at its end (Novascan Technologies, Ames, IA), which was previously calibrated by thermal tune method. 3-D piezoactuators coupled to strain gauge sensors (Physik Instrumente, Karlsruhe, Germany), allowed to place the cantilever on the region of interest with nanometric resolution and to measure the vertical displacement of the tip (z). The deflection of the cantilever (d) was measured with a quadrant photodiode (S4349, Hamamatsu, Japan) using the optical lever method. The slope of a deflection-displacement curve (d-z) obtained on a bare region of the rigid substrate was used to calibrate the relationship between cantilever deflection and photodiode signal.

Therefore, the force exerted by the cantilever was computed as F=k·d. The indentation of the sample (δ) was computed as δ = (z − z_0_) − (d − d_0_), where z_0_ and d_0_ are the positions of the contact point. To extract E, F-δ curves were analyzed by the spherical Hertz contact model (75).

Myocardial stiffness was analyzed at 7, 14 and 30 days post amputation (dpa) (n=5 per time point) and also in animals with no injury (control, n=5). For each sample time point two slices were measured. In every slice, measurements were performed in two zones: uninjured and regenerating heart. In each zone, two locations separated by ∼500µm were measured. At each of the two locations, 5 measurements were made separated by ∼10µm. E of each measurement point was the average of five force curves, obtained with ramp amplitude of 5µm and frequency of 1Hz, resulting in tip velocity of 5µm/s.

### Experimental design and statistical rationale

For LC-MS spectrometric analysis of the decellularization protocol, we used three pools of 3 native zebrafish ventricles per pool, three pools of 10 ventricles after SDS treatment per pool, and three pools of 20 decellularized ventricles per pool. A two-way ANOVA was used for comparing ECM proteins over the decellularization states. For LC-MS spectrometric analysis during regeneration, 6 ventricles per sample were pooled for 7dpa, 14dpa and 30dpa. In control samples 8 ventricles were pooled. One-way ANOVA was used to determine the statistical significant differences. Raw p-values have been controlled by adjusting the Benjamini-Hochberg False Discovery Rate (FDR).

Kruskal-Wallis followed by a Dunn’s multiple comparisons test was used to determine the qRT-PCR statistical significance of the differences at RNA level between control hearts and hearts at each regeneration time point. Mann-Whitney test was used to determine statistical significance of the differences between uninjured hearts and hearts at each time point on AFM analysis. GraphPad software was used to carry out all the statistics. All results are expressed as mean ± standard error of mean (S.E.M).

## Acknowledgements

The authors would like to thank Adriana Rodriguez-Marí and Isil Tekeli for their kind help in the initial phases of this project; and IDIBELL Clinical Proteomics Unit, Bellvitge Biomedical Research Institute (IDIBELL) for the decellularization protocol proteomic analysis. A.G.-P. was partially supported by a pre-doctoral fellowship from FPI program of the Spanish Ministry of Economy and Competitiveness (MINECO). Additional support was provided by grants from MINECO (SAF2015-69706-R), Instituto de Salud Carlos III-ISCIII/FEDER (Red de Terapia Celular - TerCel RD16/0011/0024), Generalitat de Catalunya-AGAUR (2017-SGR-899), Fundació La Marató de TV3 (201534-30), PERIS (SLT002/16/00234), and CERCA Programme / Generalitat de Catalunya. The Proteomics Laboratory at VHIO and the IDIBELL Clinical Proteomics Unit belong to ProteoRed, PRB2-ISCIII, supported by grant PT13/0001.

## Data Availability

The raw mass spectrometry data are deposited in PRIDE archive. URL: http://www.ebi.ac.uk/pride

<u>Decellularization protocol proteomic analysis</u> Project accession: PXD011627

<u>Regeneration process proteomic analysis</u> Project accession: PXD010092

## Author’s contributions

A.G.-P. and A.R. designed research; J.L.M. performed the bioinformatics analysis; A.G.-P., S.J.-D., C.G.-P., I.J., and F.C. performed research; A.G.-P. I.J., D.N., F.C., and

A.R. analyzed data; A.G.-P. and A.R. wrote the manuscript with input from all authors.

