## Supplemental Fig. 1 for "Proteomics analysis of extracellular matrix remodeling during zebrafish heart regeneration"

[8] Institució Catalana de Recerca i Estudis Avançats (ICREA).

**Running title:** Regenerating heart ECM proteome

**Keywords**: extracellular matrix; heart regeneration; proteomic analysis; atomic force microscopy.

**Supplementary Information contains:**

7 Supplementary Tables as individual sheets in attached Excel file:

**Supplementary Table 1. Proteins identified during the decellularization process.** Native zebrafish hearts, after SDS treatment (half), and at the end of decellularization (fully), were analyzed by LC-MS. Log2 raw intensities and VSN normalized intensities are given for each replicate of each sample group.

**Supplementary Table 2. Peptides identified in the decellularized samples.** Table of all the entries identified in our samples. The table contains the protein name, accession number, molecular weight and the spectral counts values obtained per each replicate.

**Supplementary Table 3. List of proteins represented with an overall of >5 spectral counts.** The table contains for each protein the gene name, the accession number, the molecular weight (MW), the ANOVA value (F), the p-value, and the adjusted p-value (FDR), the values obtained per each replicate and the average. The ECM proteins are identified with an asterisk.

**Supplementary Table 4. List of proteins represented with an overall of >5 spectral counts, normalized by VSN method.** The table contains for each protein the gene name, the accession number, the molecular weight (MW), the ANOVA value (F), the p-value, and the adjusted p-value (FDR), the values obtained per each replicate and the average. The ECM proteins are identified with an asterisk.

**Supplementary Table 5. List of proteins differentially expressed during heart regeneration.** The table contains for each protein the gene name, the accession number, the molecular weight (MW), the ANOVA value (F), the p-value, and the adjusted p-value (FDR), the values obtained per each replicate and the average. The ECM proteins are identified with an asterisk. p-value obtained from an ANOVA test of all the samples.

**Supplementary Table 6. List of the peptide identification in LC-MS analysis.**

**Supplementary Table 7. List of protein identifications.**

1 Supplementary Figure

**Supplementary Figure 1. Biological Process GO enrichment analysis on the differentially expressed proteins during cardiac regeneration.** Comparison of the enriched Biological Process GO terms on our differentially expressed proteins (**A**) and the 209 differential protein groups of Ma D. et al. (**B**).

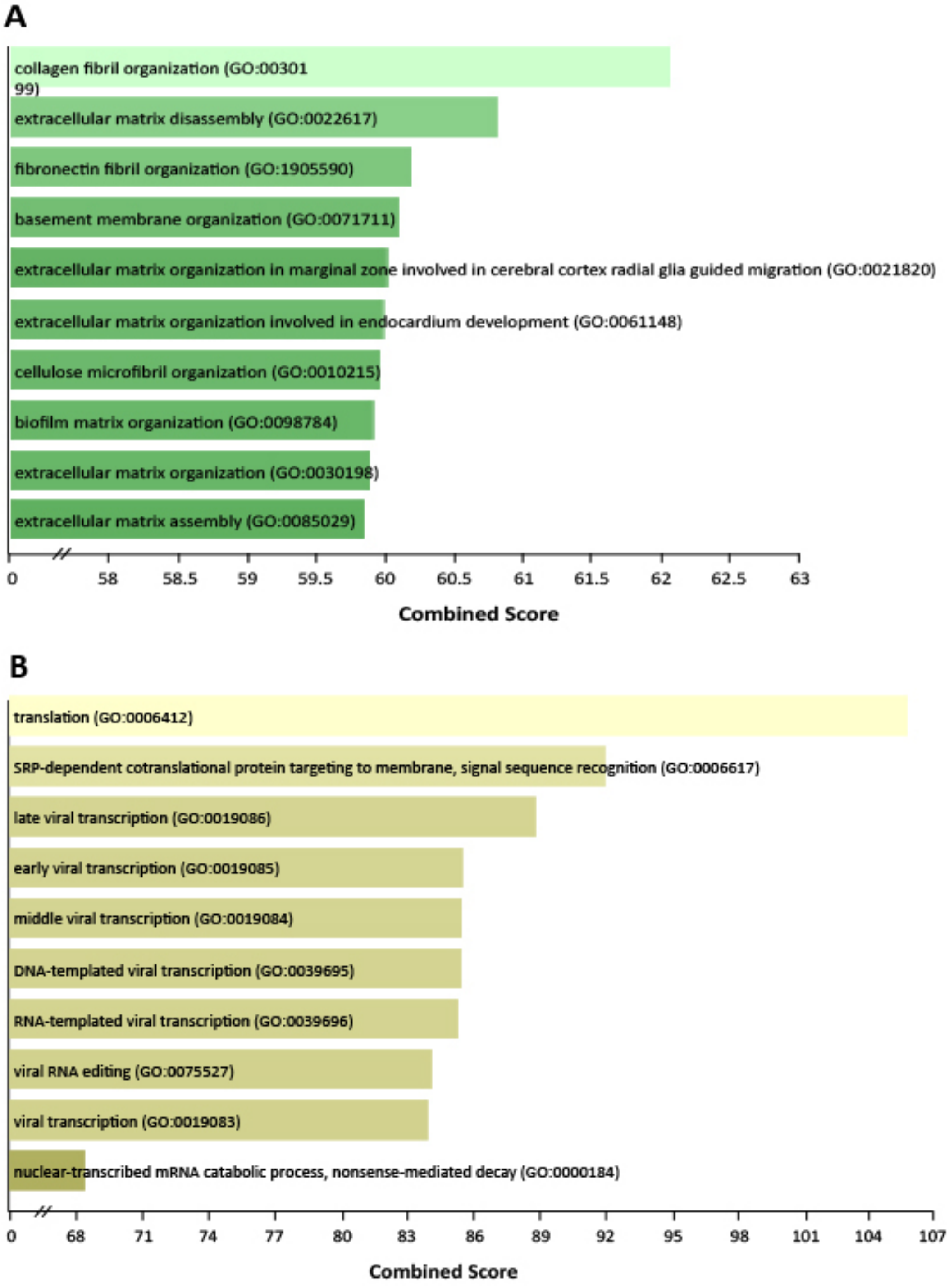


**Supplementary Fig. 1. Biological Process GO enrichment analysis on the differentially-expressed proteins during cardiac regeneration** Comparison of the enriched Biological Process GO terms on our differentially expressed proteins (A) and the 209 differential protein groups of Ma et al.[53] (B).
